# Pan-cancer genomic amplifications underlie a Wnt hyperactivation phenotype associated with stem cell-like features leading to poor prognosis

**DOI:** 10.1101/519611

**Authors:** Wai Hoong Chang, Alvina G. Lai

## Abstract

Cancer stem cells pose significant obstacles to curative treatment contributing to tumor relapse and poor prognosis. They share many signaling pathways with normal stem cells that control cell proliferation, self-renewal and cell fate determination. One of these pathways known as Wnt is frequently implicated in carcinogenesis where Wnt hyperactivation is seen in cancer stem cells. Yet, the role of conserved genomic alterations in Wnt genes driving tumor progression across multiple cancer types remains to be elucidated. In an integrated pan-cancer study involving 21 cancers and 18,484 patients, we identified a core Wnt signature of 16 genes that showed high frequency of somatic amplifications linked to increased transcript expression. The signature successfully predicted overall survival rates in six cancer cohorts (n=3,050): bladder (P=0.011), colon (P=0.013), head and neck (P=0.026), pan-kidney (P<0.0001), clear cell renal cell (P<0.0001) and stomach (P=0.032). Receiver operating characteristic analyses revealed that the performance of the 16-Wnt-gene signature was superior to tumor staging benchmarks in all six cohorts and multivariate Cox regression analyses confirmed that the signature was an independent predictor of overall survival. In bladder and renal cancer, high risk patients as predicted by the Wnt signature had more hypoxic tumors and a combined model uniting tumor hypoxia and Wnt hyperactivation resulted in further increased death risks. Patients with hyperactive Wnt signaling had molecular features associated with stemness and epithelial-to-mesenchymal transition. Our study confirmed that genomic amplification underpinning pan-cancer Wnt hyperactivation and transcriptional changes associated with molecular footprints of cancer stem cells lead to increased death risks.

**List of Abbreviations:**

TCGA: The Cancer Genome Atlas
KEGG: Kyoto Encyclopedia of Genes and Genomes
GO: Gene Ontology
ROC: Receiver operating characteristic
AUC: Area under the curve
HR: Hazard ratio
TNM: Tumor, node and metastasis
HIF: Hypoxia inducible factor
TF: Transcription factor
EMT: Epithelial-to-mesenchymal transition

## Introduction

There is a requirement for tumor cells to self-renew and proliferate in order to perpetuate tumorigenesis. It is perhaps not surprising that tumor-initiating cells or cancer stem cells share similar signal transduction processes with normal stem cells^1,2^. The ability for self-renewal and differentiation in both stem cells and cancer stem cells have converged on a common pathway known as Wnt signaling^3,4^. Wnt proteins are highly conserved across the animal kingdom, functioning as developmentally important molecules controlling cell fate specification, cell polarity and homeostatic self-renewal processes in embryonic and adult stem cells^5^. Wnts are a group of glycoproteins serving as ligands for the frizzled receptor to initiate signaling cascades in both canonical and non-canonical pathways^6^. Beyond embryogenesis, Wnt proteins control cell fate determination in adults where they regulate homeostatic selfrenewal of intestinal crypts and growth plates^7-9^.

Wnt signaling is the product of an evolutionary adaptation to growth control in multicellular organisms, and it has now become clear that aberrations in this pathway contributes to deranged cell growth associated with many disease pathologies including cancer^10^. Loss-of-function mutations in genes that inhibit the Wnt pathway lead to ligand-independent constitutive activation of Wnt signaling in hepatocellular carcinoma^11^, colorectal cancer^12^, gastric cancer^13^ and acute myeloid leukemia^14^. Thus, inhibition of Wnt signaling would hold great promise as therapeutic targets^15^. A small molecule inhibitor ICG-001 functions to inhibit the degradation of the Wnt repressor Axin and treatment of colon cancer cell lines with this inhibitor resulted in increased apoptosis^16^. Antibodies against Wnts and frizzled receptors have also demonstrated antitumor effects^17,18^.

Much of the previous research on Wnt genes and cancer have focused on somatic mutations and transcriptional dysregulation of Wnt pathway members. Activating mutations of β-catenin have been implicated in adrenocortical tumorigenesis^19^ and multiple gastrointestinal cancers^20^. Downregulation of a Wnt antagonist *DKK1,* a downstream target of β-catenin, is also observed in colorectal cancer^21^. However, there is limited understanding on the role of somatic copy number alterations in Wnt pathway genes as well as their downstream targets on driving tumor progression and patient prognosis. Studies examining the transcriptional dysregulation of Wnt pathway genes offered limited insights into whether differences in transcript abundance were caused by genomic amplifications or losses.

Given the complexity of Wnt signaling in cancer, it is important to investigate genomic alterations alongside transcriptional regulation of *all* genes associated with Wnt signaling in a comparative approach. We hypothesize that pan-cancer transcriptional aberrations in Wnt signaling is caused by genomic amplifications of a group of genes known as Wnt drivers and that transcriptional profiles of driver genes are important predictors of patient outcome. We conducted a pan-cancer analysis on 147 Wnt signaling genes, which involved positive and negative regulators of the pathway alongside their downstream targets. We analyzed 18,484 matched genomic and transcriptomic profiles representing 21 cancer types to determine whether *1)* somatic copy number amplifications are drivers of hyperactive Wnt signaling, *2)* Wnt driver genes harbor clinically relevant prognostic information and *3)* crosstalk exists between Wnt driver genes, tumor hypoxia and signaling pathways associated with stem cell function. We demonstrate that overexpression of Wnt driver genes resulted in significantly poorer survival outcomes in six cancer types involving 3,050 patients. Hyperactivation of Wnt signaling is linked to loss of cell adhesion and molecular features of stemness. Overall, our findings would facilitate the development of improved therapies through the inhibition of Wnt driver genes in a stratified manner.

## Materials and Methods

A total of 147 genes associated with active and inactive Wnt signaling were retrieved from the Kyoto Encyclopedia of Genes and Genomes (KEGG) database listed in Table S1.

### Study cohorts

Genomic and transcriptomic profiles of 21 cancers were generated by The Cancer Genome Atlas (TCGA) initiative^22^ (n=18,484) (Table S2). For transcriptomic profiles, we retrieved Illumina HiSeq rnaseqv2 Level 3 RSEM normalized data from the Broad Institute GDAC Firehose website. For somatic copy number alterations analyses, we retrieved GISTIC datasets^23^ using the RTCGAToolbox package to access Firehose Level 4 copy number variation data. Level 4 clinical data were retrieved using RTCGAToolbox for survival analyses.

### Somatic copy number alterations analyses

GISTIC gene-level table provided discrete amplification and deletion indicators for all tumor samples. Amplified genes were denoted as positive numbers: ‘1’ represents amplification above the threshold or low-level gain (1 extra copy) while ‘2’ represents high-level amplification (2 or more extra copies). Deletions were denoted as negative values: ‘-1’ represents heterozygous deletion while ‘-2’ represents homozygous deletion.

### Determining the 16-gene scores and hypoxia scores

16-Wnt-gene scores for each patient were determined from the mean log2 expression values of 16 genes: *WNT2, WNT3, WNT3A, WNT10B, FZD2, FZD6, FZD10, DVL3, WISP1, TBL1XR1, RUVBL1, MYC, CCND1, CAMK2B, RAC3* and *PRKCG.* Hypoxia scores were computed from the mean log2 expression values of 52 hypoxia signature genes^24^. For analyses in Figures 5 and 7, patients were separated into four groups using median 16-gene scores and median hypoxia scores or median *EZH2* expression values as thresholds. Nonparametric Spearman’s rank-order correlation tests were employed to investigate the relationship between 16-gene scores and hypoxia scores or *EZH2* expression values.

### Differential expression analyses

To compare Wnt gene expression between tumor and non-tumor samples, gene expression profiles for both sample types were separated into two files based on TCGA barcode information. RSEM expression values were converted to log2(x + 1) scale. To compare changes in gene expression between high- and low-score groups, patients were median dichotomized based on their 16-gene scores in each cancer type. Differential expression analyses were performed using the R limma package employing the linear model and Bayes method. P value adjustments were conducted using the Benjamini-Hochberg false discovery rate method.

### Biological enrichment and transcription factor analyses

To ascertain which biological pathways and signaling processes were significantly enriched as a result of Wnt hyperactivation, differentially expressed genes obtained from comparing high- and low-score patients were mapped against the KEGG and Gene ontology (GO) databases using GeneCodis^25^. Differentially expressed genes were also mapped against the Reactome database^26^. The Enrichr tool was used to determine whether differentially expressed genes were enriched with binding targets of stem cell-associated transcription factors^27,28^. Genes were mapped against the ChEA and ENCODE databases using Enrichr.

### Survival analysis

The R survminer and survival packages were used for Kaplan-Meier and Cox proportional hazards regression analyses to determine if the expression levels of the 16 signature genes were significantly associated with overall survival. The ability of the 16-gene signature to predict overall survival when used in combination with hypoxia scores or *EZH2* expression levels was also examined. Univariate Cox regression analyses were performed on each of the individual 16 genes in 20 cancer types (where survival information is available) to determine the contribution of each gene in predicting overall survival. Univariate analyses were also performed on the gene set as a signature (by taking the mean expression scores of the 16 genes) to determine its ability in predicting overall survival. Multivariate Cox regression analyses were employed to demonstrate the independence of the signature to tumor staging parameters. Hazard ratios (HR) and confidence intervals were determined from Cox models where HR greater than one (P<0.05) indicated that a covariate was positively associated with even probability (increased hazard) and negatively linked to survival length. The non-significant relationship between scaled Schoenfeld residuals and time supported the proportional hazards assumption; this was tested using the R survival package. Kaplan-Meier analyses were employed to confirm results obtained from Cox regression. Patients were first median-separated into low- and high-score groups based on the expression of the 16 genes (detailed above) for Kaplan-Meier analyses. Statistical difference between high- and low-score patient groups was evaluated using the log-rank test. Receiver operating characteristic analyses were performed using the R survcomp package to assess the predictive performance (sensitivity and specificity) of the signature in relation to tumor stage. Area under the ROC curves (AUCs) were calculated using survcomp. AUC values can fall between 1 (perfect marker) and 0.5 (uninformative marker).

All plots were generated using ggplot2 and pheatmap packages implemented in R^29^. The InteractiVenn tool^30^ was employed to generate the Venn diagram in Figure S2.

## Results

### Pan-cancer genomic alterations of Wnt signaling lead to dysregulated transcriptional response in tumors

A list of 147 genes involved in the Wnt signal transduction pathway was retrieved from the KEGG database (Table S1). They include genes in both canonical and non-canonical Wnt pathways along with their downstream targets. A literature search was conducted to manually curate these genes into two categories: 1) genes associated with active Wnt signaling (90 genes) and 2) genes associated with repressed Wnt signaling (50 genes) (Fig. 1A). To systematically evaluate the extent of Wnt dysregulation across cancers, we analyzed genomic and transcriptomic datasets from 18,484 patients representing 21 cancer types^22^. To determine whether genomic alterations were present in the 147 genes, we evaluated the frequency of somatic copy number alterations across all 21 cancers.

**Figure 1.**
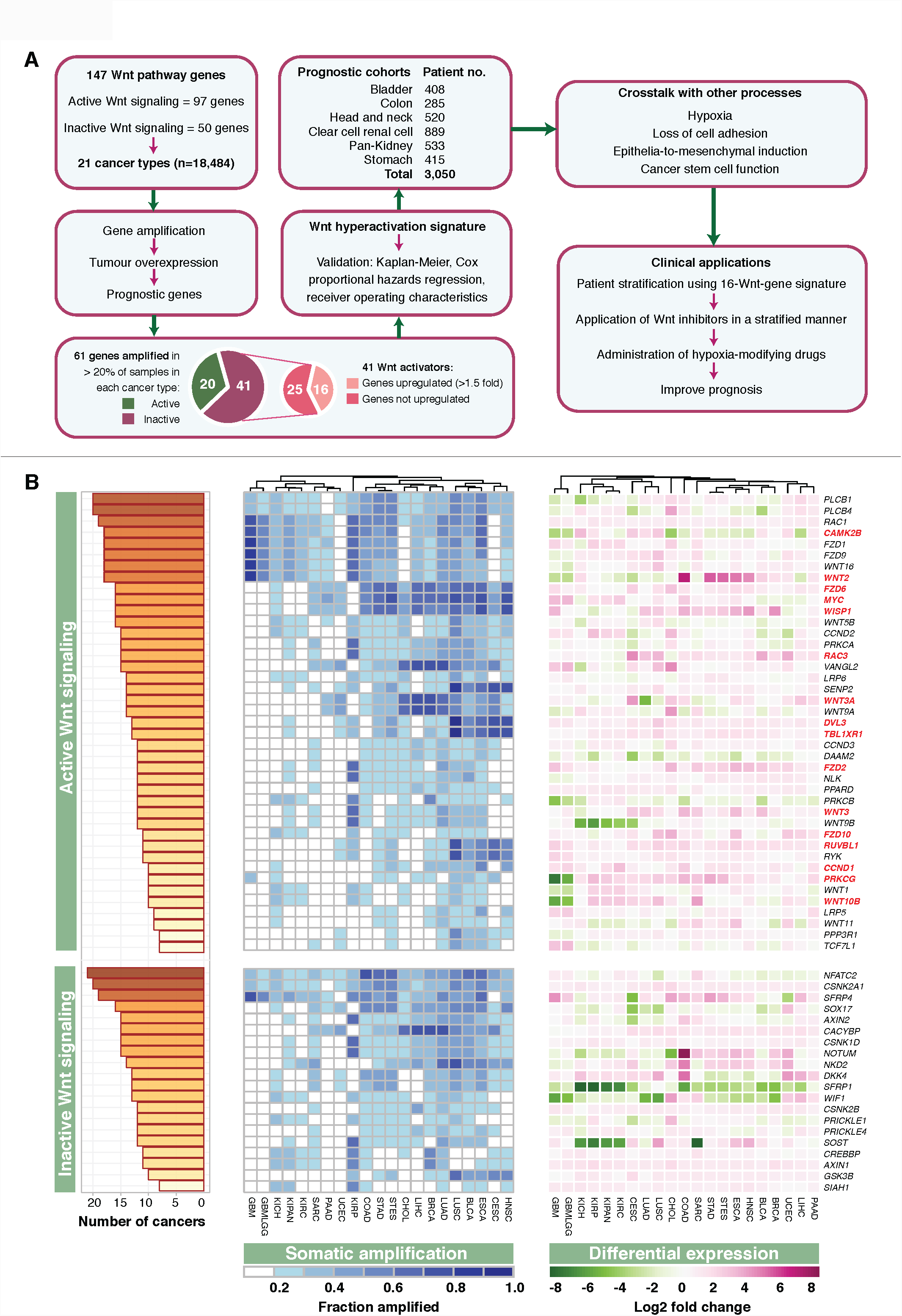
Pan-cancer core drivers of Wnt signaling. **(A)** Schematic diagram depicting the study design and the identification of core Wnt driver genes subsequently representing the 16-gene signature. A total of 147 Wnt signaling genes representing both canonical and non-canonical pathways alongside their downstream targets were obtained from the KEGG database. Genes were grouped into two categories depending on whether they were associated with active or inactive Wnt signaling. Somatic copy number variations in all 147 genes were determined in 21 cancer types. A total of 61 genes were recurrently amplified in at least 20% of tumors in each cancer type. They included 41 genes associated with active Wnt signaling. Of the 41 genes, 16 genes (core Wnt drivers) were upregulated in tumor compared to non-tumor samples in at least 8 cancer types. Cox proportional hazards regression and Kaplan-Meier analyses were performed using the 16-gene signature, which demonstrated its ability to predict overall survival in at least six cancer types: bladder, colon, head and neck, clear cell renal cell, papillary renal cell, chromophobe renal cell and stomach cancers (n=3,050). Associations of the 16-Wnt-gene signature with cancer stem cell features, tumor hypoxia and cell adhesion were investigated. Potential clinical applications of the signature were proposed. (B) Somatic amplification and differential expression profiles of 61 Wnt genes. Cumulative bar chart depicts the number of cancer types with at least 20% of tumors with somatic gains. The heatmap on the left shows the extent of genomic amplifications for each of the 61 genes separated into ‘active’ and ‘inactive’ Wnt signaling categories across 21 cancer types. Heatmap intensities indicate the fraction of the cohort in which a given gene is gained or amplified. The columns were ordered using hierarchical clustering with Euclidean distance metric to reveal cancers that have similar somatic amplification profiles. The heatmap on the right demonstrates differential expression values (log2) between tumor and non-tumor samples for each of the 61 genes. Genes marked in red represent the 16 Wnt driver genes. These are genes that were amplified in at least 20% of tumors in at least 8 cancers and genes that were overexpressed (fold-change > 1.5) in at least 8 cancers. Refer to Table S2 for cancer abbreviations.

Focusing on genomic amplifications that occurred in at least 20% of samples in each cancer type and amplification events that were present in at least one-third of cancer types (> 8 cancers), we observed that 61 genes were recurrently amplified (Fig. 1B). Of these 61 genes, 41 genes were associated with active Wnt signaling while 20 genes were linked to repressed Wnt signaling (Fig. 1A). Some of the most amplified genes found in at least 95% of cancer types included genes from both canonical (*FZD1, FZD9, WNT16, WNT2, SFRP4, CSNK2A1* and *RAC1)* and non-canonical Wnt pathways (*PLCB1, PLCB4, CAMK2B* and *NFATC2)* (Fig. 1B).

When comparing the frequency of Wnt gene amplifications between cancers, interesting associations were observed. Cancers that affect organ systems working together to perform a common function, i.e. gastrointestinal tract, exhibited similar patterns of genomic amplifications where most of the 61 genes were amplified in at least 20% of tumors. Hierarchical clustering on amplification frequencies using Euclidean distance metric revealed that gastrointestinal cancers of the colon (COAD), stomach (STAD), bile duct (CHOL) and liver (LIHC) were clustered together, implying that there was a significant degree of conservation in genetic aberration of Wnt signaling in these cancers (Fig. 1B). In contrast, cancers of the brain and central nervous system (GBMLGG and GBM) had the least number of amplified genes; 11 and 12 genes respectively (Fig. 1B).

We reason that somatic amplification events that were linked with transcriptional overexpression could represent candidate Wnt drivers, given that positive correlation between RNA and DNA levels would imply a gain of function. We performed differential expression analyses on the 90 genes involved in active Wnt signaling (Table S1) using tumor and non-tumor samples from each cancer type (Table S2). We observed that 28 genes were overexpressed (fold change > 1.5) in at least 8 or more cancers. Of the 28 genes, we identified 16 genes that were also recurrently amplified (Fig. 1A, B). These 16 genes were prioritized as core Wnt driver candidates representative of multiple tumors: *WNT2, WNT3, WNT3A, WNT10B, FZD2, FZD6, FZD10, DVL3, WISP1, TBL1XR1, RUVBL1, MYC, CCND1, CAMK2B, RAC3* and *PRKCG* (Fig. 1B).

### Pan-cancer prognostic relevance of the newly identified core Wnt drivers

We rationalize that the gain of function of the core Wnt drivers could influence patient outcome. Univariate Cox proportional hazards regression analyses were performed on the transcriptional profiles of each of the 16 Wnt drivers on 20 cancers where survival information is available. A vast majority of the core Wnt driver genes were significantly associated with poor prognosis (hazard ratio [HR] above 1, P<0.05) (Fig. S1). Interestingly, there were variations in the number of prognostic genes between cancers. Esophageal cancer (ESCA) had no prognostic genes and only two genes were prognostic in sarcoma (SARC) and cholangiocarcinoma (CHOL). In contrast, clear cell renal cell carcinoma (KIRC) and the pan-kidney cohort (KIPAN) involving chromophobe renal cell, papillary renal cell and clear cell renal cell carcinoma had 13 and 10 prognostic genes respectively (Fig. S1). To determine whether core Wnt driver genes harbored prognostic information as a gene set, we calculated expression scores for each patient in each cancer type by taking the mean expression of the 16 Wnt drivers. Patients were subsequently median-dichotomized into low- and high-score groups for survival analyses. Remarkably, when the core Wnt drivers were considered as a gene signature, we observed that patients with high scores had significantly poorer survival rates in six cancer cohorts (n=3,050): bladder (P=0.011), colon (P=0.013), head and neck (P=0.026), pan-kidney (P<0.0001), clear cell renal cell (P<0.0001) and stomach (P=0.032) (Fig. 2).

**Figure 2.**
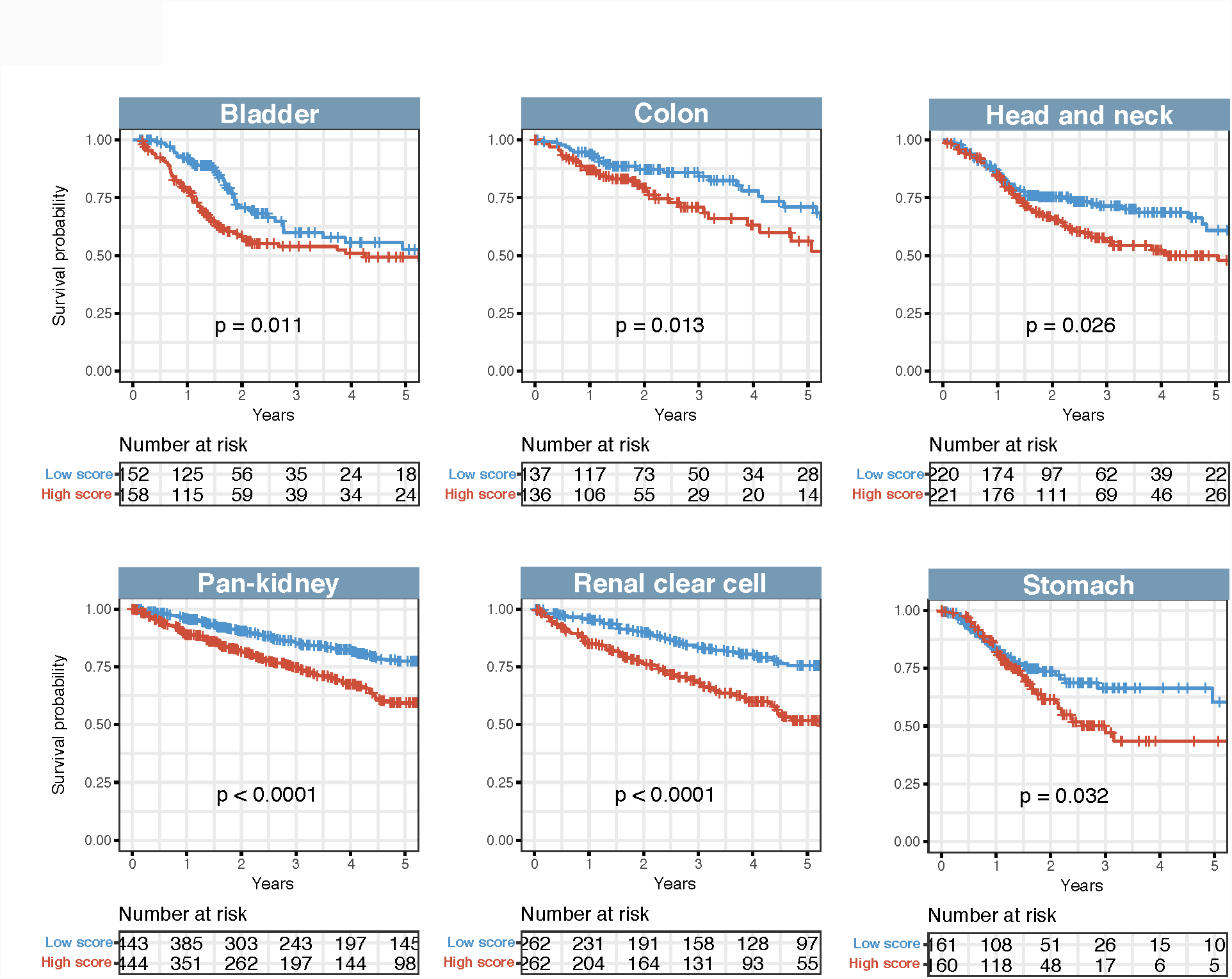
Survival analyses using the 16-Wnt-gene signature in six cancer cohorts. Kaplan-Meier analyses of overall survival on patients stratified into high- and low-score groups using the 16-gene signature. P values were determined from the log-rank test.

To determine whether the 16-Wnt-gene signature harbored independent prognostic value over current tumor, node and metastasis (TNM) staging system, the signature was evaluated on patients grouped according to tumor stage; early (stages 1 and/or 2), intermediate (stages 2 and/or 3) and late (stages 3 and/or 4). Patients were first separated by tumor stage followed by median-stratification based on their 16-gene scores into low- and high-score groups within each stage category. Regardless of tumor stage, the signature retained its predictive value where high-score patients consistently had higher risk of death: early stage (bladder: P=0.0043, colon: P=0.03, head and neck: P=0.024, pan-kidney: P=0.045, clear cell renal cell: P=0.0008 and stomach: P=0.036), intermediate stage (colon: P=0.029, pan-kidney: P=0.012, clear cell renal cell: P=0.00031 and stomach: P=0.028) and late stage (pan-kidney: P=0.0014, clear cell renal cell: P=0.00032 (Fig. 3). Taken together, this suggests that another level of patient stratification beyond that of TNM staging is afforded by the 16-gene signature, especially for patients with early stage cancer where tumors are more heterogeneous.

**Figure 3.**
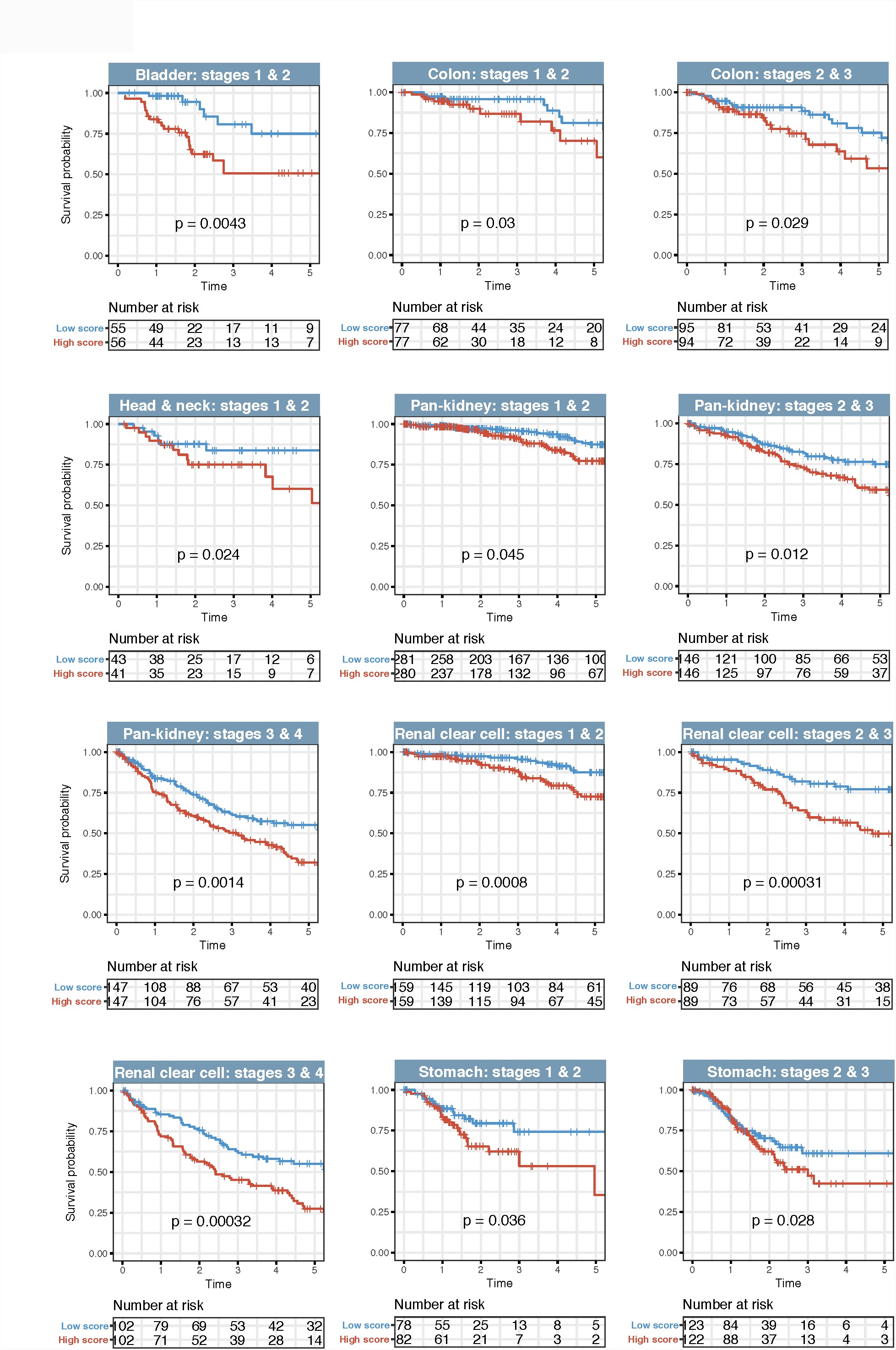
The 16-Wnt-gene signature is independent of TNM stage. Kaplan-Meier analyses were performed on patients categorized according to tumor TNM stages that were further stratified using the 16-gene signature. The signature successfully identified patients at higher risk of death in all TNM stages. P values were determined from the log-rank test. TNM: tumor, node, metastasis.

Multivariate Cox regression analyses were performed to further confirm that the 16-Wnt-gene signature was independent of TNM staging. Indeed, in all six cancer types, the signature remained prognostic when controlling for TNM stage (Table S3). High-score patients had significantly higher risk of death even when TNM stage was taken into account: bladder (HR=1.409, P=0.015), colon (HR=1.561, P=0.018), head and neck (HR=1.378, P=0.036), pan-kidney (HR=1.738, P<0.0001), clear cell renal cell (HR=2.146, P<0.0001) and stomach (HR=1.457, P=0.035) (Table S3).

We next employed the receiver operating characteristic (ROC) method to assess the predictive performance (specificity and sensitivity) of the 16-gene signature in determining 5-year overall survival rates. As revealed by the area under the ROC curves (AUCs), we confirmed that the signature had consistently outperformed TNM staging in all six cancers: bladder (AUC=0.707 vs. AUC=0.626), colon (AUC=0.673 vs. AUC=0.652), head and neck (AUC=0.624 vs. AUC=0.606), pan-kidney (AUC=0.779 vs. AUC=0.717), clear cell renal cell (AUC=0.740 vs. AUC=0.717) and stomach (AUC=0.754 vs. AUC=0.561) (Fig. 4). Importantly, when the signature was used as a combined model with TNM staging, we observed a further increase in AUC suggesting that the signature offered incremental predictive value: bladder (AUC=0.713), colon (AUC=0.723), head and neck (AUC=0.663), pan-kidney (AUC=0.833), clear cell renal cell (AUC=0.818) and stomach (AUC=0.757) (Fig. 4).

**Figure 4.**
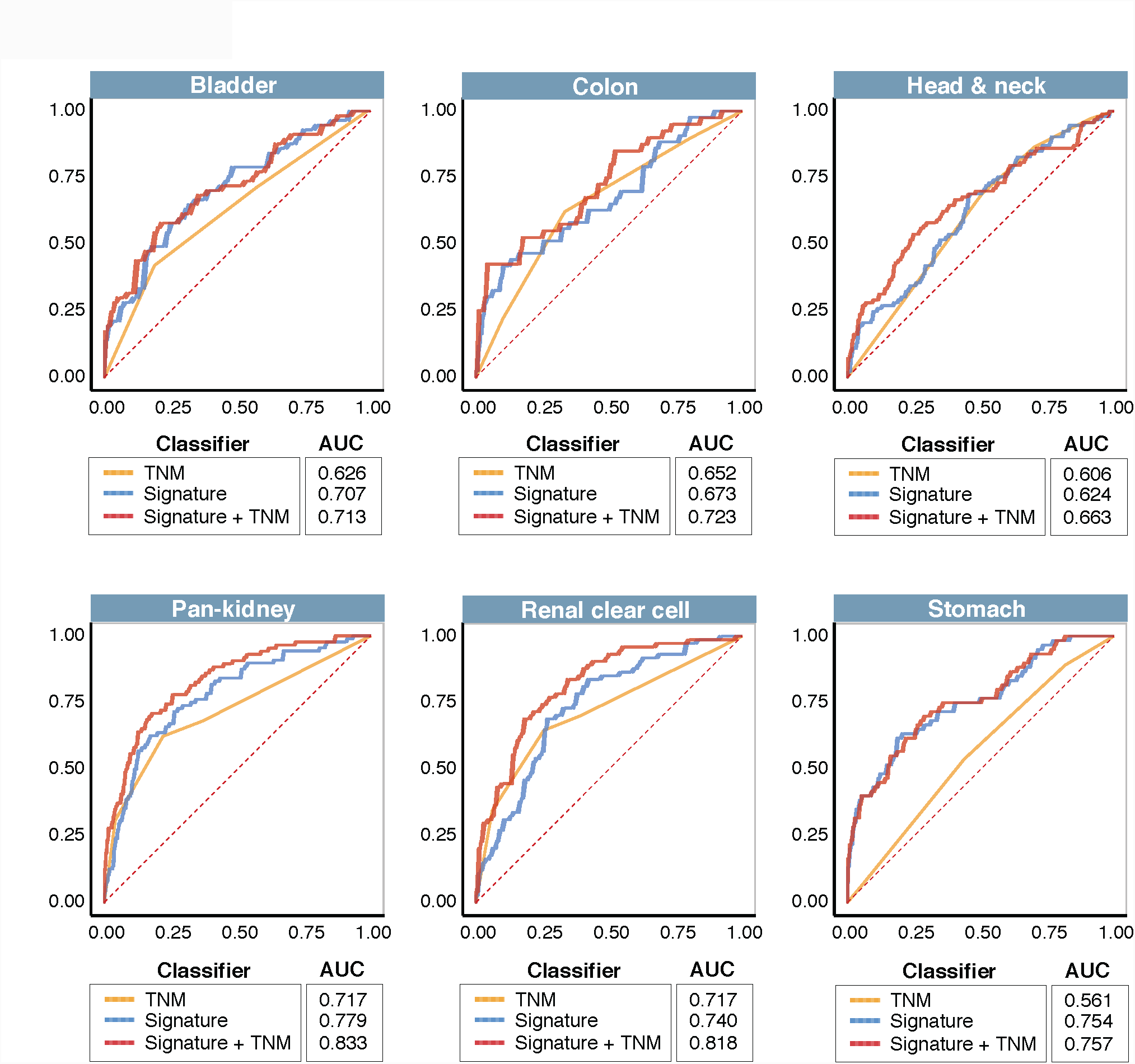
Predictive performance of the 16-Wnt-gene signature is superior to TNM staging. Prediction of five-year overall survival was assessed using the receiver operating characteristic (ROC) analysis to determine specificity and sensitivity of the signature. ROC curves were generated based on the 16-gene signature, TNM stage and a combination of the signature and TNM stage. AUC: area under the curve. TNM: tumor, node, metastasis. TNM staging were in accordance with previous publications employing TCGA datasets^33,34^.

### Association of Wnt drivers with tumor hypoxia

Poor vascularization in solid tumors results in tumor hypoxia that is frequently associated with very poor prognosis due to reduced effectiveness of chemotherapy and radiotherapy^31^. Furthermore, the stabilization of the hypoxia inducible factor (HIF) in hypoxic tumor microenvironments can promote metastasis and cancer progression leading to poor prognosis^32-34^. An emerging view on cancer stem cells postulates that hypoxic regions could serve as stem cell niches to provide an oxidative DNA damage-buffered zone for cancer stem cells^35,36^. Moreover, crosstalk between HIFs and stem cell signal transduction pathways (Wnt, Notch and *Oct4)* have been reported^37,38^. For instance, HIF-1α can interact with (β-catenin to promote stem cell adaptation in hypoxic conditions^39^.

Multiple evidence suggests that Wnt signaling may be influenced by the extent of hypoxia within the tumor microenvironment. We reason that hypoxia could further enhance Wnt signaling to allow cancer stem cells to persist, which together contribute to even poorer survival outcomes in patients. Integrating hypoxia information with the 16-Wnt-gene signature would enable the evaluation of the crosstalk between both pathways and its clinical relevance. We predict that patients with more hypoxic tumors would have higher expression of Wnt driver genes, which may imply that these patients have higher proportions of tumor-initiating cells with hyperactive Wnt signaling. To assess tumor hypoxia levels, we utilized a computationally derived hypoxia gene signature comprising of 52 genes^24^. Hypoxia scores were calculated for each patient as the average expression of the 52 genes. Interestingly, significant positive correlations were observed between the 16-Wnt-gene scores and hypoxia scores in bladder (rho=0.365, P<0.0001) and clear cell renal cell cancers (rho=0.305, P<0.0001), suggesting that in these two cancers, hypoxic tumors had higher expression of core Wnt drivers (Fig. 5A).

**Figure 5.**
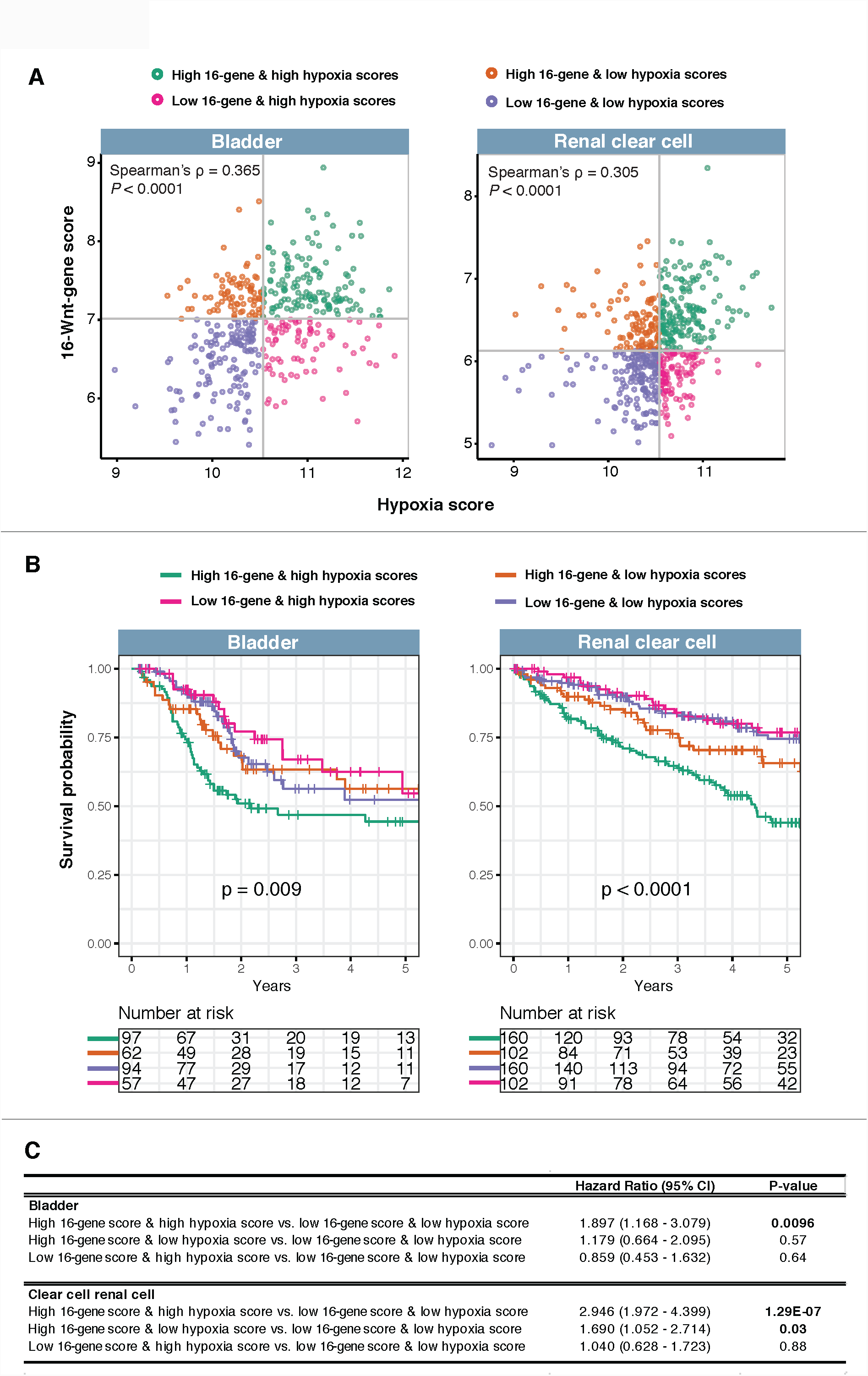
Positive associations between the 16-gene signature and tumor hypoxia in bladder and clear cell renal cell cancers. **(A)** Scatter plots show significant positive correlation between 16-gene scores and hypoxia scores as determined by Spearman’s rank-order correlation analyses. Patients were separated and color-coded into four categories based on median 16-gene and hypoxia scores. (**B)** Kaplan-Meier analyses were performed on the four patient categories to determine the effects of the combined relationship between hypoxia and the Wnt signature on overall survival. (**C)** Univariate Cox proportional hazards analysis of the relation between the 16-gene signature and hypoxia. CI: confidence interval.

To determine the clinical relevance of this positive association, we separated patients into four groups: 1) high scores for both 16-gene and hypoxia, 2) high 16-gene score and low hypoxia score, 3) low 16-gene score and high hypoxia score and 4) low scores for both 16-gene and hypoxia (Fig. 5A). Kaplan-Meier analyses were performed on the four patient groups and we observed that the combined relation of Wnt hyperactivation and hypoxia was significantly associated with overall survival in both cancers: bladder (P=0.009) and clear cell renal cell (P<0.0001) (Fig. 5B). Notably, patients with high hypoxia and high 16-gene scores had significantly higher mortality rates compared to those with low hypoxia and low 16-gene scores: bladder (HR=1.897, P=0.0096) and clear cell renal cell (HR=2.946, P<0.0001) (Fig. 5C). Overall, our results suggest that the joint effect of elevated hypoxia and Wnt signaling is linked to more aggressive disease states.

### Wnt hyperactivation is responsible for epithelial-to-mesenchymal transition properties through decreased cell adhesion

Given the poor survival outcomes in patients with high 16-gene scores, we wanted to assess the biological consequences of hyperactive Wnt signaling. Patients were median-stratified into two categories, high- and low-score, for differential expression analyses. For each cancer, the number of differentially expressed genes (−1 > log2 fold-change > 1, P<0.05) were 1,543 (bladder), 1,164 (colon), 984 (head and neck), 659 (pan-kidney), 943 (clear cell renal cell) and 328 (stomach) (Table S4) (Fig. S2). Gene ontology (GO) enrichment analyses revealed enrichment of biological processes consistent with those of cancer stem cells: cell proliferation, cell differentiation, embryo development and cell morphogenesis (Fig. 6A). Moreover, despite their diverse tissue origins, high-score patients from all six cancers exhibited remarkably similar biological alterations (Fig. 6A) (Table S4). For example, high-score patients appear to show a phenotype associated with loss of cell adhesion properties. Genes involved in regulating cell adhesion were downregulated and the ‘cell adhesion’ GO term was among the most enriched ontologies across all six cancers (Fig. 6A). As a further confirmation, differentially expressed genes were mapped to the KEGG database and enrichments of ontology related to cell adhesion molecules were similarly observed (Fig. 6B). A third database known as Reactome^26^ was used in functional enrichment analyses. Comparing results from both KEGG and Reactome analyses revealed enrichments of additional processes related to oncogenesis and Wnt signaling; e.g. altered metabolism, PPAR signaling, MAPK signaling, TGF-β signaling, Hedgehog signaling, calcium signaling, collagen synthesis and degradation, focal adhesion and chemokine signaling (Fig. 6B, C). Within the tumor microenvironment, collagen can modulate extracellular matrix conformation that could paradoxically promote tumor progression^40,41^. Indeed, we observed the enrichment of numerous collagen-related Reactome pathways: assembly of collagen fibrils, collagen biosynthesis, collagen formation, collagen chain trimerization and collagen degradation (Fig. 6C). Overall, our results suggest that elevated mortality risks in high-score patients could potentially be due to loss of cell adhesion and aggravated disease states exacerbated by Wnt hyperactivation.

**Figure 6.**
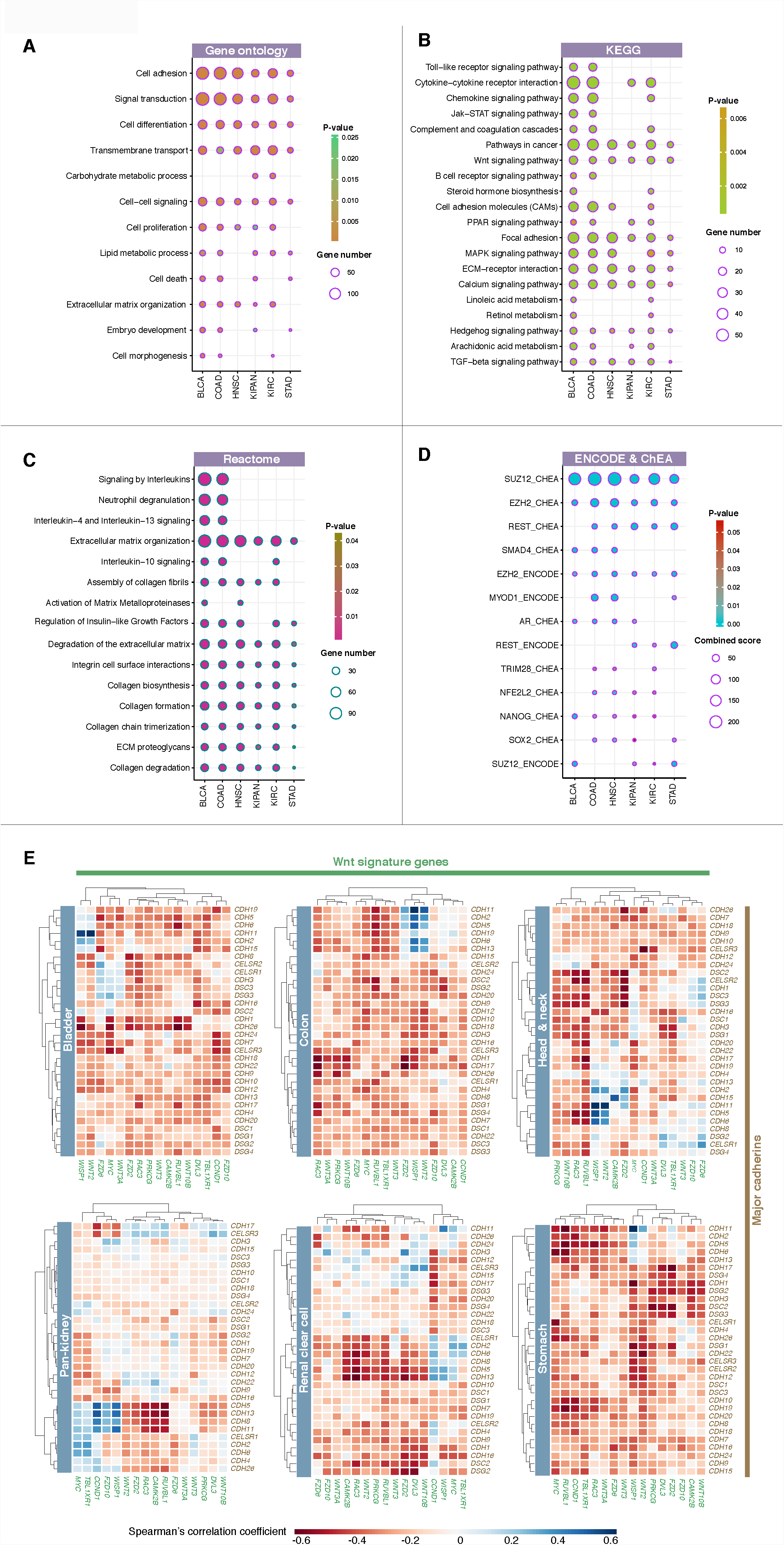
Wnt hyperactivation is associated with a cancer stem cell-like phenotype. Patients were median separated into high- and low-score groups using the 16-gene signature for differential expression analyses. Enrichments of biological processes on differentially expressed genes were determined by mapping the genes to (**A)** Gene Ontology, (**B)** KEGG and (C) Reactome databases. Significantly enriched pathways or ontologies for all six cancer cohorts were depicted. (**D)** Differentially expressed genes were enriched for targets of stem cell-related transcription factors (Nanog, Sox2, Smad4, EZH2 and SUZ12) as confirmed by mapping to ENCODE and ChEA databases. Refer to Table S2 for cancer abbreviations. (**E)** Significant negative correlations between the expression profiles of individual Wnt driver genes and 32 major cadherin genes. Heatmaps were generated based on Spearman’s correlation coefficient values.

To determine the extent of the loss of adhesive properties in tumor cells expressing high levels of Wnt driver genes, we examined the expression profiles of 32 genes from the major cadherin superfamily. Major cadherins are a group of highly conserved proteins that encode at least five cadherin repeats, which include type I and II classical cadherins (*CDH1, CDH2, CDH3, CDH4, CDH5, CDH6, CDH7, CDH8, CDH9, CDH10, CDH11, CDH12, CDH13, CDH15, CDH18, CDH19, CDH20, CDH22, CDH24* and *CDH26*), 7D cadherins (*CDH16* and *CDH17*), desmosomal cadherins (*DSC1, DSC2, DSC3, DSG1, DSG2, DSG3* and *DSG4)* and CELSR cadherins (*CELSR1, CELSR2* and *CELSR3)*^*42*^. Spearman’s correlation analyses between major cadherins and each of the individual Wnt driver genes revealed that the 16 genes exhibited a global pattern of negative correlation with major cadherins across all six cancer types (Fig. 6E). Taken together, these results provide further support to the notion on loss of cadherin-mediated cell adhesion in tumor cells with hyperactive Wnt signaling, which may act in concert to promote neoplastic progression.

### A role for *EZH2* histone methyltransferase in cancer stem cells

When analyzing transcription factor (TF) binding to differentially expressed genes described in the previous section, we observed that these genes were enriched for targets of several notable TFs such as EZH2, SUZ12, Nanog, Sox2 and Smad4 (Fig. 6D). Sox2 and Nanog are well-known stem cell markers^43^ while EZH2 and SUZ12 are part of the polycomb repressive complex 2 responsible for epigenetic regulation during embryonic development^44,45^ (Fig. 6D). The enrichment of target genes of these TFs supports the hypothesis that Wnt hyperactivation is associated with cancer stem cell properties. Aberrations in *EZH2* and *SUZ12* have been linked to cancer progression^46-50^ and overexpression of *EZH2* is associated with poor prognosis^51^. Direct crosstalk between *EZH2* function and Wnt signaling has been reported where *EZH2* was shown to inhibit Wnt pathway antagonists to activate Wnt/β-catenin signaling leading to increased cellular proliferation^52^. Moreover, *EZH2* inhibits E-cadherin expression via lncRNA H19 to promote bladder cancer metastasis^53^.

Since EZH2 binding targets were enriched among differentially expressed genes (confirmed by both ChEA and ENCODE databases) and given the role of EZH2 in cell adhesion and Wnt signaling, we reason that *EZH2* would be overexpressed in tumors with hyperactive Wnt signaling. Indeed, significant positive correlations were observed between 16-Wnt-gene scores and *EZH2* expression in renal cancers: pan-kidney (rho=0.203, P<0.0001) and clear cell renal cell (rho=0.233, P<0.0001) (Fig. 7A). Patients were further grouped by their 16-gene scores and *EZH2* expression profiles into four categories: 1) high 16-gene score and high *EZH2* expression, 2) high 16-gene score and low *EZH2* expression, 3) low 16-gene score and high *EZH2* expression and 4) low 16-gene score and low *EZH2* expression (Fig. 7A). Interestingly, patients with high 16-gene score that concurrently had high *EZH2* expression had the poorest survival outcomes compared to the others: pan-kidney (P<0.0001) and clear cell renal cell (P<0.0001) (Fig. 7B). This suggests that Wnt hyperactivation and *EZH2* overexpression could synergize to drive tumor progression resulting in significantly higher death risks: pan-kidney (HR=3.444, P<0.0001) and clear cell renal cell (HR=3.633, P<0.0001) (Fig. 7C).

**Figure 7.**
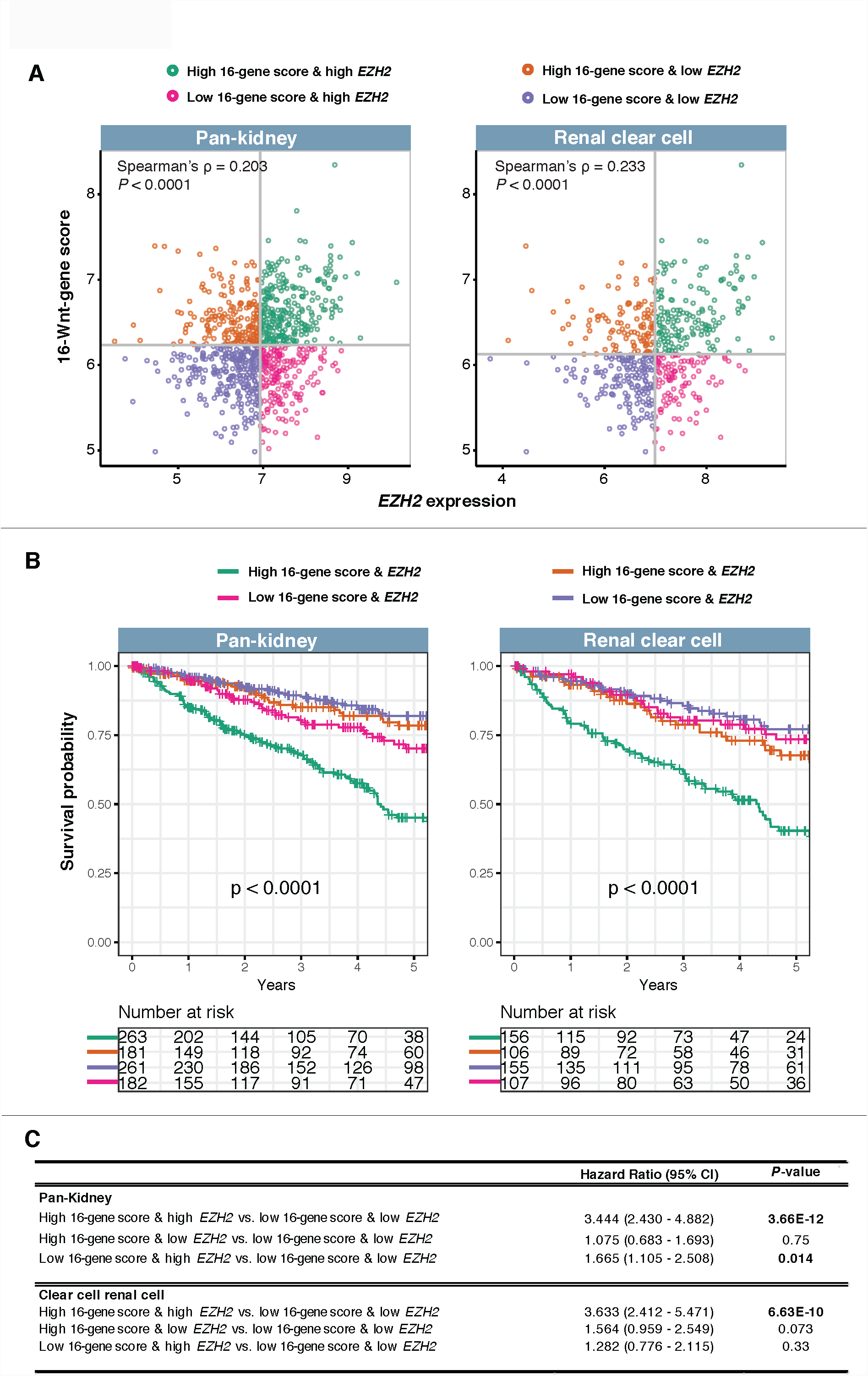
Positive associations between the 16-gene signature and *EZH2* expression in renal cancers. **(A)** Scatter plots show significant positive correlation between 16-gene scores and *EZH2* expression as determined by Spearman’s rank-order correlation analyses. Patients were separated and color-coded into four categories based on median 16-gene score and *EZH2* expression. (**B)** Kapla n-Meier analyses were performed on the four patient categories to determine the effects of the combined relationship between *EZH2* expression and the Wnt signature on overall survival. (**C)** Univariate Cox proportional hazards analysis of the relation between the 16-gene signature and *EZH2* expression. CI: confidence interval.

## Discussion and Conclusion

We performed a comprehensive pan-cancer analysis of 147 Wnt pathway genes in 18,484 patients from 21 different cancer types to unravel the intricacies of Wnt regulation of cancer phenotypes. Taking into account genomic, transcriptomic and clinical data, we demonstrated that overexpression of Wnt genes is underpinned by somatically acquired gene amplifications (Fig. 1). We found that differential Wnt activation contributed to significant heterogeneity in disease progression and survival outcomes. Focusing on 16 core Wnt drivers that were recurrently amplified and overexpressed, our results confirmed that Wnt hyperactivation drove malignant progression that is conserved across diverse cancer types (Fig. 2, 3, 4). Our newly developed 16-Wnt-gene signature could predict patients with more aggressive disease states who may benefit from treatment with small molecule inhibitors of Wnt^16,54,55^.

Copy number amplification and concomitant overexpression of WNT driver genes in bladder, colon, head and neck, renal and stomach cancers were significantly associated with stem cell-like molecular features (Fig. 6). The transcriptional profiles of 16 Wnt drivers were negatively correlated with the expression of a vast majority of major cadherin genes involved cell adhesion; a process that may drive epithelial-to-mesenchymal transition (EMT)^56^(Fig. 6E). This is consistent with the role of Wnts as inducers of EMT^57^. Patients with high expression of Wnt driver genes exhibited enriched biological processes involving cytokine, TGF-β and Hedgehog signaling (Fig. 6); these components are also implicated in regulating EMT induction^57^. TGF-β activation orchestrates signaling events activating downstream effectors such as Smad proteins that play essential roles in cellular differentiation^58^. Indeed, we observed that dysregulated genes in tumors with hyperactive Wnt signaling were enriched for Smad4 targets (Fig. 6D). Smads can bind to Zeb proteins to repress E-cadherin expression during the onset of EMT^59,60^. The downregulation of major cadherins in tumors expressing high levels of Wnt drivers (Fig. 6E) could thus be a combined result of aberrant Wnt and TGF-β signaling.

Patients with Wnt hyperactivation exhibited additional molecular features of undifferentiated cancer stem cells. We observed enrichments of stem cell-related TFs such as Nanog, Sox2 and polycomb proteins (SUZ12 and EZH2) as upstream targets of Wnt-associated dysregulated genes; this pattern was consistent across the different cancer types (Fig. 6D). Patients with Wnt hyperactivation phenotypes could have poorly differentiated tumors reminiscent of cancer stem cells given their preferential misexpression of genes normally associated with embryonic stem cell function (Fig. 7). The distinction between cancer stem cells and normal stem cells is of paramount interest. Molecular footprints of stemness identified from analyzing the transcriptional changes between high- and low-16-WNT-gene-score patients could provide additional evidence of cancer stem cell identity in these tumors that is linked to poor overall prognosis.

Our results also demonstrated that Wnt signaling is positively correlated with tumor hypoxia in bladder and clear cell renal cell cancers. Patients with more hypoxic tumors had higher 16-Wnt-gene scores, suggesting that tumor hypoxia may contribute to the activation of Wnt genes. These patients could benefit from the use of hypoxia-modifying drugs such as carbogen and nicotinamide shown to be effective in bladder cancer^61^ to reduce tumor hypoxia, which may consequently dampen Wnt signaling. Crosstalk between Wnt signaling and hypoxia has been demonstrated in multiple cancers. (β-catenin expression is induced by hypoxia in liver cancer, which contributes to increased EMT, invasion and metastasis^62^. Overexpression of HIF-1α promoted invasive potential of prostate cancer cells through β-catenin induction, while the silencing of β-catenin in HIF-1α expressing cells resulted in increased and reduced epithelial marker and mesenchymal marker expression respectively^63^. Hypoxia-induced EMT is further enhanced by the addition of recombinant Wnt3a or is repressed by inhibiting β-catenin^64^. Indeed, our results confirmed that increased expression of Wnt driver genes was associated with a global downregulation of major cadherin genes consistent across six cancer types, which may occur through hypoxia-mediated processes (Fig. 6E). We observed that in clear cell renal cell carcinoma, patients with more hypoxic tumors who also had higher Wnt signature scores concomitant with a 2.9-fold higher risk of death (Fig. 5C). Interestingly, renal cancers have a high incidence of VHL mutations^65^. VHL is a protein involved in proteasomal degradation of HIF-1α ^66^. VHL antagonizes the Wnt pathway through β-catenin inhibition in renal tumors^67^, meaning that VHL mutations would derepress Wnt signaling and create a pseudohypoxic environment to further promote the expression of Wnt pathway genes. Our results will open up new research avenues for investigating the role of the 16 Wnt drivers and potential crosstalk with VHL-mediated HIF signaling in renal cancer.

In summary, we identified Wnt pathway genes that were recurrently amplified and overexpressed across 21 diverse cancer types. A core set of 16 genes known as Wnt drivers were preferentially expressed in high-grade tumors linking to poor overall survival. This signature is a prognostic indicator in six cancer types involving 3,050 patients and is independent and superior to tumor staging parameters, providing additional resolution for patient stratification within similarly staged tumors. We demonstrated clinically relevant relationships between the 16-gene signature, cancer stem cells, cell adhesion, tumor hypoxia and *EZH2* expression. Hence, aggressive tumor behavior and survival outcomes are, in part, driven by Wnt hyperactivation. Furthermore, we reported evidence for crosstalk between Wnt signaling and other embryonic stem cell pathways (TGF-α signaling, Nanog, Sox2 and polycomb repressive complex 2) confirming that these pathways do not operate in isolation and that interactions between them could add to the complexity of neoplastic progression. Prospective validation in clinical trials and additional functional studies on individual Wnt drivers are needed before they can be harnessed for therapeutic intervention.

## Authors contribution

WHC and AGL designed the study, analyzed the data and interpreted the data. AGL supervised the research. WHC and AGL wrote the initial manuscript draft. AGL revised the manuscript draft and approved the final version.

## Supplementary figures and tables

**Figure S1. Prognosis of each of the 16 signature genes in 20 cancer types as determined using Cox regression analyses.** Both columns (cancer types) and rows (Wnt genes) were ordered using hierarchical clustering (Euclidean distance metric). Grey boxes represent non-prognostic genes. Heatmap intensities represent hazard ratios of prognostic genes that were significant (P<0.05).

**Figure S2. Venn diagram depicts a six-way comparison of the differentially expressed genes identified from high-score versus low-score patients in all six cancer cohorts.** Numbers in parentheses represent the number of differentially expressed genes (−1 > log2 fold-change > 1, P<0.05) in each cancer.

**Table S1.** List of 147 genes associated with Wnt signaling.

**Table S2.** Abbreviations and number of tumor and non-tumor samples in TCGA cancers.

**Table S3.** Univariate and multivariate Cox proportional hazards analysis of risk factors associated with overall survival in multiple cancers.

**Table S4.** Differentially expressed genes between high- and low 16-Wnt-score patient groups in six cancers.

